# Ensemble-based genome-scale modeling predicts metabolic differences between macrophage subtypes in colorectal cancer

**DOI:** 10.1101/2023.03.09.532000

**Authors:** Patrick E. Gelbach, Stacey D. Finley

## Abstract

Colorectal cancer (CRC) shows high incidence and mortality, partly due to the tumor microenvironment, which is viewed as an active promoter of disease progression. Macrophages are among the most abundant cells in the tumor microenvironment. These immune cells are generally categorized as M1, with inflammatory and anti-cancer properties, or M2, which promote tumor proliferation and survival. Although the M1/M2 subclassification scheme is strongly influenced by metabolism, the metabolic divergence between the subtypes remains poorly understood. Therefore, we generated a suite of computational models that characterize the M1- and M2-specific metabolic states. Our models show key differences between the M1 and M2 metabolic networks and capabilities. We leverage the models to identify metabolic perturbations that cause the metabolic state of M2 macrophages to more closely resemble M1 cells. Overall, this work increases understanding of macrophage metabolism in CRC and elucidates strategies to promote the metabolic state of anti-tumor macrophages.

## 2 Introduction

Colorectal cancer is the fourth most common cancer in the world, and the second most common cause of cancer-related death in the United States^1^. Even with the current standard of care and new therapies, CRC patients have a high rate of relapse, and resistance to therapy is a key contributor to morbidity and mortality. Part of the difficulty in effectively treating CRC is the complexity of the tumor microenvironment (TME), which arises from the diverse range of cell types and extracellular matrix components surrounding the tumor. The TME is composed of cancer cells, stromal cells such as fibroblasts and immune cells, and extracellular matrix components such as collagen and proteoglycans^2–4^. The interactions between these different cell types and matrix components can influence the behavior of cancer cells and affect response to therapy^5, 6^. Moreover, these interactions can promote an immunosuppressive environment and support drug resistance. Thus, understanding the interactions between the different cell types and matrix components in the TME is crucial for developing effective therapies for CRC.

One aspect of the TME that is not fully understood is its metabolic profile. Tumor cells have a distinct metabolic state compared to normal cells, characterized by high rates of glucose uptake and lactate production. This metabolic shift, known as the Warburg effect, allows cancer cells to generate energy and biomass at an accelerated rate, which supports tumor growth and progression^7, 8^. The cells surrounding the tumor influence and are influenced by the cancer cells, and thus experience metabolic changes and the emergence of phenotypically distinct subgroups within a given cell type^9^. Macrophages are among the most common cells in the TME, and are responsible for a wide variety of immune activity in the body; therefore, the distinct cell subtypes within the macrophage population are of particular interest^10, 11^. Simplistically, macrophages are classically considered to be polarized into two broad categories: a pro-inflammatory (M1) state, or a pro-resolving (anti-inflammatory, M2) condition. The M1 state is responsible for initiating and sustaining immune responses. M1 cells are often activated in response to foreign infections, causing the secretion of cytokines and other bactericidal mediators. The M2 state is associated with a reduction in microenvironment inflammation, as the cells release anti-inflammatory mediators and collagen, which encourage tissue repair^12, 13^. More recent evidence shows that macrophages can exist in a continuum of states; however, consideration of the pro- and anti-tumor classification is well-supported and historically accepted, and therefore is deserving of further research^14, 15^.

The clear distinction in cellular behavior of macrophages has for a long time been viewed as strongly related to differences in intracellular metabolism^16–18^. Technological advances in omics-level profiling provide insights into the metabolic preferences of macrophages. Metabolomics, the comprehensive and systematic analysis of intracellular metabolites, enables characterization of the cell’s metabolic status^19, 20^. Metabolomics studies show that M1 cells demonstrate dependence on glycolysis and the catabolism of arginine to nitric oxide. In contrast, M2 cells are shown to preferentially utilize oxidative phosphorylation and are oriented towards the production of urea and polyamines, which are used as mediators of wound healing^13, 21, 22^.

However, most metabolomic studies of macrophage subtypes are limited in their scope, as they focus on quantifying the levels of intracellular metabolites. Although it is useful to quantify the metabolite levels, it is difficult to infer cell function and to assess metabolic state of a cell solely based on the levels of individual metabolites. Rather, the rates of the complex network of biochemical reactions that the metabolites participate in are highly indicative of the metabolic state of a cell^23^. However, since it is difficult to capture flux measurements at the whole-cell level, most studies have focused on central metabolism and only a few accessory pathways^24–27^.

Beyond technical limitations, there are context-dependent effects on macrophage metabolism that are difficult to capture. For example, although it is known that cancer cells encourage macrophages to convert from an M1 to an M2 state (also known as tumor associated macrophages, or TAMs), a quantitative understanding of the effects of cancer cell-induced metabolic reprogramming of macrophages is relatively unknown^28^. It is not clear what metabolic alterations exist in cancer-induced macrophages. With a better understanding of the metabolic states of macrophages in the tumor microenvironment, it may be possible to identify strategies to modulate macrophage metabolism and improve the outlook for CRC patients.

Genome-scale metabolic models (GEMs) are promising tools to address the limitations of purely experimental metabolomics-based studies. These models consist of two connected matrices: a stoichiometric (S) matrix showing all the cell’s known metabolites and metabolic reactions, and a gene-protein-reaction (GPR) rules matrix showing the enzymes and genes known to be linked to those reactions. The S-matrix allows for the study of cellular metabolism with linear algebra approaches (such as flux balance analysis)^29, 30^. The GPR rules permit multi-omic data integration, allowing the GEM to act as a scaffold onto which collected data can be overlayed, thus generating context-specific models^31^. It is therefore possible to predict the distribution of material (flux) through the metabolic network to quantitatively characterize cell state and cell phenotype. The predicted flux values are constrained by known biological properties (such as thermodynamic limits or measured cell activity) or by context-specific confines (such as the availability of extracellular nutrients). Thus, genome-scale modeling is often called a “constraint-based” analysis of metabolism^32^. The modeling technique emerged in studies of bacterial and yeast metabolism but has been increasingly utilized in the field of cancer biology. This is because genome-scale modeling takes advantage of -omics datasets to infer function and phenotype among all metabolic reactions without a dependence on difficult-to-obtain kinetic parameters that are required for other modeling techniques^33, 34^.

There has been limited work in using genome-scale modeling to understand macrophage biology and phenotypic divergence. Bordbar *et al.* studied the host-pathogen interactions of human alveolar macrophages in tuberculosis, but did not account for variation in M1 and M2 states^35^. In another paper, the same group modeled M1 and M2 distinction and macrophage activation in a murine leukemia cell line^36^. However, the authors did not account for differences in structure of the metabolic model between the two macrophage subgroups, instead simulating the same model with different cellular goals. In this work, we generate the first example of human-specific models of cancer-associated M1 and M2 macrophage metabolism, based on *in vivo*-omics data. We analyze the structure of the metabolic models and predicted fluxes to reveal differences in the metabolic states of M1 and M2 macrophages in CRC. The metabolites and reactions shown in our models to distinguish M1 and M2 cells match the commonly accepted M1 and M2 metabolic markers, providing confidence that the models can be used to further investigate macrophage metabolism. Thus, we apply the models to identify therapeutic targets to induce M2 cells towards an M1-like state, potentially informing future immunotherapies in colorectal cancer.

## 3 Materials and Methods

### 3.1 RNA Sequencing Data

Li *et al.* collected 1,591 cells from 11 patients with primary stage CRC and profiled the cells using a single-cell RNA sequencing protocol^37^. Eum *et al.* processed the data collected by Li *et al.*, identified CRC-associated macrophages, and categorized the cells as M1 and M2 based on the presence of accepted cell surface markers and by comparing to reference transcriptomes^38^.

In total, 98 M1 macrophages and 56 M2 macrophages were identified and analyzed with single-cell RNA sequencing. This sequencing analysis identified 3,216 and 3,187 measured genes in M1 and M2 macrophages, respectively, that are present in the Recon3D model of human metabolism^39^. We pooled the measured single-cell data into subtype-specific “pseudo-bulk” transcriptomics profiles to use for development of the macrophage GEMs^40^.

### 3.2 Data Integration

We integrated the RNA sequencing data into the Recon3D model of human metabolism using the constraint-based reconstruction and analysis (COBRA) Toolbox (version 3.0, implemented in MATLAB [Mathworks, Inc.])^41^. Recon3D is among the most recent and complete models for human metabolism, consisting of 3,288 open reading frames, 13,543 reactions and 4,140 metabolites in 103 distinct pathways in a generic human cell. Several algorithms exist to integrate transcriptomics data into a generic model of species metabolism, thus allowing for the generation of context-specific models that can be applied to infer cellular phenotype and intracellular flux distributions^42^. Each integration approach has its own set of steps and settings used to determine which reactions should be maintained in or removed from the base model, given the transcriptomics data. In order to minimize the effect of algorithm selection on cellular predictions, we selected five commonly used algorithms and applied each one with the aforementioned transcriptomics data. Specifically, we use: (1) integrative metabolic analysis tool (iMAT), (2) gene inactivity moderated by metabolism and expression (GIMME), (3) cost optimization reaction dependency assessment (CORDA), (4) integrative network inference for tissues (INIT), and (5) FASTCORE^43–47^. We describe each, and its relevant properties and parameters, below.

#### 3.2.1 iMAT

This method first divides genes profiled in the transcriptomics data into low-, moderate-, or high-expression levels. iMAT then maximizes the presence of reactions corresponding to highly expressed genes and minimizes the presence of reactions linked to low expression genes. The method thus finds the optimal tradeoff between retaining reactions related to highly expressed genes and removing reactions related to genes with low expression. We used the top 25% of expressed genes as the cutoff for “high” expression, and the bottom 25% as “low” expression.

#### 3.2.2 GIMME

This method removes “inactive” reactions. These are reactions where the corresponding RNA transcript level is below a specific lower threshold or reactions that are not required for user-defined core functionality. GIMME requires an objective function to be maximized, and we used ATP maintenance. To ensure that the choice of objective function did not largely impact the generation of model, we also set biomass maintenance as a selected objective. However, there was no difference in the models generated.

#### 3.2.3 CORDA

The CORDA method determines the high-, medium-, and low-confidence reactions based on the transcriptomics data. The method then includes all high-confidence reactions and aims to maximize inclusion of high confidence reactions while minimizing inclusion of low-confidence reactions. We define the confidence level based on gene expression: “high” corresponds to the genes with expression in the top 25%, “medium” confidence reactions as middle 50%, and “low” as bottom 25% expression levels.

#### 3.2.4 INIT

INIT assigns weights to each reaction based on the corresponding gene’s expression level, then finds the optimal tradeoff between keeping reactions with high weights and removing reactions with low weights. We calculated weights as:

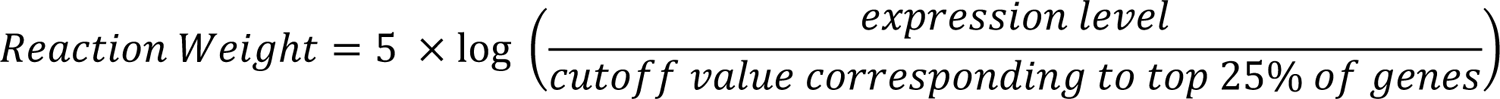

#### 3.2.5 FASTCORE

This method first defines core reactions: reactions corresponding to genes with high expression. FASTCORE then searches for a flux consistent network, one that has a nonzero flux for all of the core reactions and a minimum number of additional reactions. We define genes with high expression as those in the top 25% expression level of the transcriptomics data.

#### 3.2.6 Consistency Between Algorithms

We maintained consistency between algorithms by using the same definition of high and low gene expression, set of important metabolic tasks relevant to macrophages, and objective functions. As described above, when there was a threshold required for a gene to be considered highly expressed, we used the top 25% of the measured data. Similarly, if a cutoff was required for a gene to be designated as low expression, we used the bottom 25% of measured genes. We ensured inclusion of reactions and metabolites responsible for set of metabolic tasks for which there is clear evidence in macrophages (**Supplementary Table 1**). We also ensured the inclusion of synthetic or pseudo-reactions that represent cellular maintenance functions and are commonly used as objective functions when simulating the model. Namely, we maintained the biomass reaction, biomass maintenance reaction, ATP maintenance reaction, and ATP maximization reactions. Because those reactions represent a lumped biological function and are not directly measurable in transcriptomics, they would otherwise be removed by the integration algorithms.

### 3.3 Consensus Models Generated via REDGEM Pipeline and Benchmark Analysis

In addition to the individual M1 and M2 models produced by each of the five integration approaches, we produced a single consensus model for M1 macrophages and a single model for M2 macrophages using the REDGEM pipeline. REDGEM provides a minimal model from a user-provided set of reactions of interest^48^. We applied the approach to the list of consensus model components for each subtype (the genes, reactions, and metabolites found in three or more of the five derived models), thus obtaining a consolidated model that contained the high-confidence components. We compare the consensus-derived models of M1 and M2 macrophage metabolism produced by REDGEM to identify similarities and differences in the genes, reactions, and subsystems for the two macrophage subtypes. We then analyzed all 12 context-specific GEMs (10 obtained from integrating transcriptomics data and two consensus models obtained from REDGEM) using MEMOTE^49^. The MEMOTE test suite analyzes a GEM for proper annotation and formal correctness, benchmarking the model across four domains: (1) *annotation* – ensuring the model is annotated according to community standards; (2) *basic tests* – checking the correctness of the model, including metabolite formulas, charge, and GPR rules; (3) *biomass* – confirming that the model can produce required biomass precursors and thus simulate cell growth; and (4) *stoichiometry* – reporting stoichiometric errors and permanently blocked reactions.

### 3.4 Flux Predictions

#### 3.4.1 Flux Balance Analysis

We performed flux balance analysis (FBA) to calculate the steady state reaction fluxes that account for the stoichiometric and mass balance constraints given by the S-matrix, together with the flux bounds for each reaction. As the system is underdetermined, an objective function is specified to predict the optimal set of reaction fluxes required to maximize or minimize the objective function^50^. Though a single cellular objective (i.e., maximizing biomass or minimizing ATP usage) is often reasonable when simulating prokaryotic metabolism, it may not be applicable for higher-order species. In particular, multiple objective simulations may better represent the metabolic behavior of mammalian cells^51–53^. Thus, we selected objective functions that collectively comprise the cellular goals of macrophages. For M1 cells, we selected biomass maintenance, ATP maintenance, and activity of the inducible nitric oxide synthase (iNOS) enzyme, which is shown to be important for M1 macrophages^54, 55^. For M2 cells, we used biomass maintenance, ATP maintenance, and activity of the arginase 1 (ARG1) enzyme, as it has been used to identify M2 cells^13, 56, 57^.

#### 3.4.2 Flux Sampling

We performed flux sampling to explore the achievable flux distribution of context-specific GEMs using the Riemannian Hamiltonian Monte Carlo (RHMC) algorithm^58^. Flux sampling does not require an objective function and can produce many sets of feasible flux distributions, thus providing confidence values and insight into the range of possible fluxes for each reaction^59^. In total, we generated 50,000 flux samples per model subtype.

### 3.5 Comparison of Reaction Flux Distributions

With distributions containing a range of flux values for each reaction, we compared the activity of each shared metabolic reaction present in the M1 and M2 models using the Kullback-Leibler (KL) divergence^60^. The KL divergence is a statistical metric categorizing the difference between two distributions. A KL divergence of 0 indicates that the two distributions are equivalent. Previous work has categorized reaction flux distributions as being in “close agreement” if the KL divergence is less than 0.05, “medium agreement” if between 0.05 and 0.5, and “largely divergent” if greater than 0.5^61, 62^. Thus, we applied the same cutoffs when evaluating the KL divergence for the flux distribution for each reaction present in M1 and M2 macrophage GEMs. It is important to note that the KL divergence metric is one-sided. That is, the KL divergence of M1 versus M2 is not equivalent to the KL divergence of M2 versus M1. For that reason, we performed the divergence calculation in each direction and then averaged the two divergence values to arrive at a single distance metric for each reaction. Similarly, we calculated the average KL divergence metric for each subsystem present in the M1 and M2 models to determine the pathways whose flux values differ between the M1 and M2 models.

### 3.6 PageRank Analysis

We performed a flux-weighted PageRank analysis to determine the relative importance of the metabolic reactions present in both the M1 and M2 GEMs^63–65^. This approach provides a graph theory-based measure of connectivity of the metabolic network, scaled by the flux through the reaction. The PageRank algorithm was originally developed to determine the most important nodes for a set of search engine results, but has since been applied to other graph-based questions, including understanding the connectivity of metabolic networks. The flux-weighted PageRank value indicates how significant each metabolic reaction is, allowing for efficient comparison between two GEMs and identification of reactions that likely drive the observed metabolic phenotype.

### 3.7 Reaction Knockouts

We simulated a reaction knockout by setting the upper and lower bounds for the flux through that reaction to zero. We subsequently performed flux sampling to predict the reaction fluxes in the knockout metabolic model. We were particularly interested in how the reaction knockout affects the flux distributions for all reactions originally classified as highly divergent between the unperturbed M1 and M2 models. Thus, we used t-distributed stochastic neighbor embedding (t-SNE), an unsupervised dimensionality reduction technique, to represent the distributions for the highly divergent reactions^66^. In doing so, we represented the predicted flux distributions for the unperturbed and knockout models as points in two-dimensional space. We calculated the cosine distance between points for the unperturbed M1 model and knockout models. That distance was normalized to the distance between the unperturbed M1 and M2 models. Finally, the percent change was calculated as:

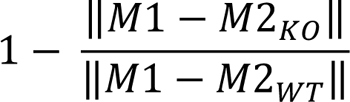

## 4 Results

### 4.1 Ensemble modeling of macrophage metabolism

We generated *in silico* models of macrophage metabolic activity by integrating data into a generic GEM of human metabolism (Recon3D). Using patient-derived data (described in the *Materials and Methods* section), we employed five methods (CORDA, GIMME, IMAT, INIT, and FASTCORE) to integrate the patient-derived transcriptomics data and produce context-specific metabolic models. Each approach uses distinct algorithms to maintain or remove metabolic reactions from a generic GEM based on gene expression data. By using five approaches, we generated a suite of five context-specific metabolic models for each of the macrophage subtypes (M1 and M2). We maintained consistency in input parameters (the same base model, data, and identical ranges of the top and bottom 25% of measured genes designated as “highly expressed” and “lowly expressed”, respectively). However, there was significant variation in size of the resulting models (**Figure 1A**). INIT produced the smallest models, while CORDA and GIMME retained most model components and thus produced large models that were similar in size to the input model. Furthermore, for a particular data integration algorithm, the size of the M1 and M2 models were nearly the same (**Figure 1B and 1C**). This reveals that the model integration approach, rather than the experimental data, was the primary cause for divergence in model composition, as shown in the pairwise clustering of M1 and M2 models.

**Figure 1:**
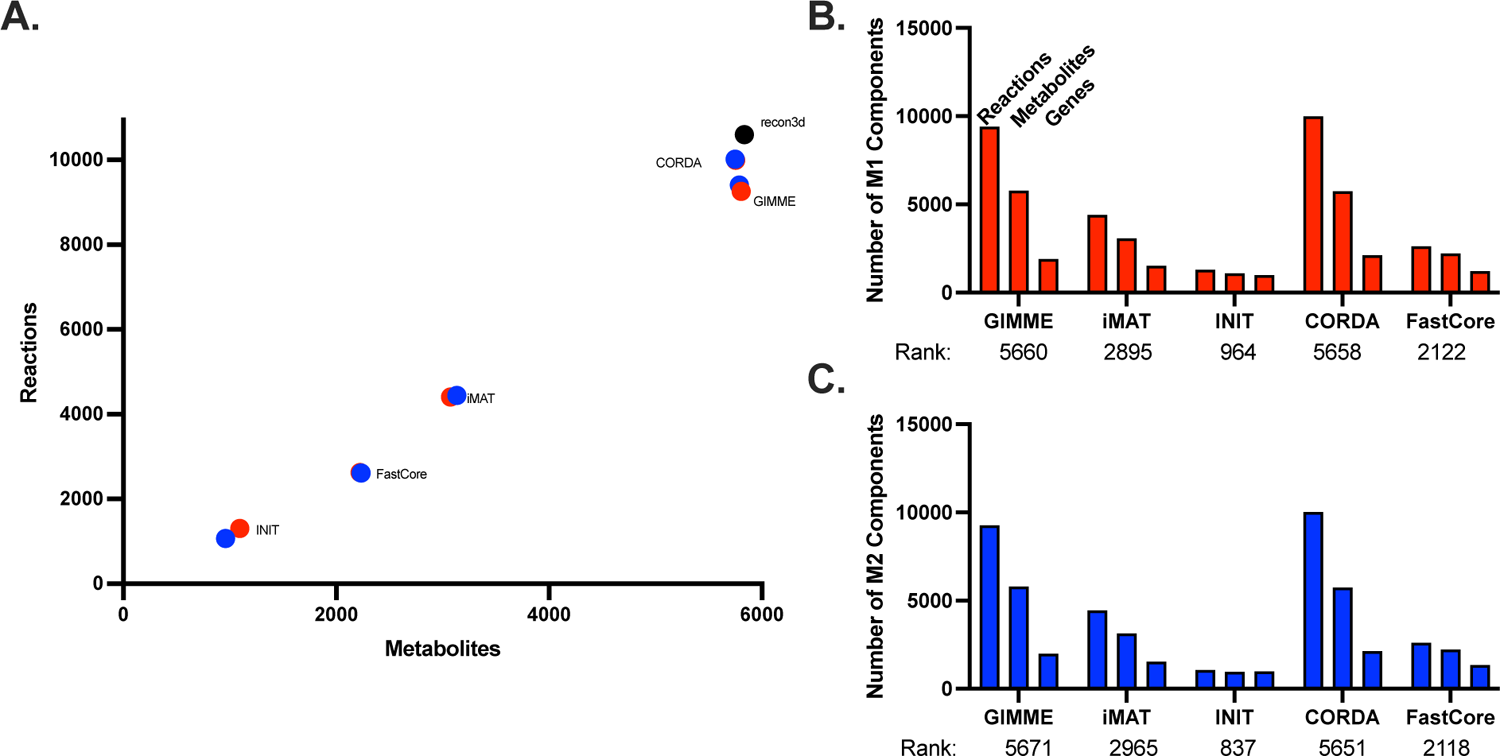
Size of context-specific genome-scale models. **A.** Size of models generated by integrating patient-derived transcriptomics data from M1 and M2 macrophages using five distinct approaches. **B.** The number of components (genes, reactions, and metabolites) and stoichiometric matrix rank for each model generated.

We then sought to analyze consistency of the suite of generated models. We first evaluated the context-specific models with the MEMOTE test suite, which assesses model feasibility and correctness (particularly, stoichiometric and thermodynamic consistency and appropriate annotation). We found no stoichiometric, charge, or thermodynamic errors. No model had orphan metabolites or dead-end reactions, and there were sufficient GPR rules for each model reaction. None of the models generated had errors, pointing to the validity of each integration approach.

We next compared model components (metabolic genes, reactions, and metabolites) across the five models generated for each subtype. We calculated the number of models that contained a particular component. That is, we determined the presence of each model gene, reaction, and metabolite across the ensembles of models. This is shown in **Figure 2A**. As expected, when the ensemble cutoff (the number of models that must contain the component) is more restrictive, fewer model components meet the threshold. We further investigated the relationship between the integration algorithm used and presence or absence of a model component, as shown in the Venn diagrams of **Figure 2B**. Because INIT tends to produce smaller models, most model components included by that approach are also found in the other techniques. Conversely, the models produced by CORDA and GIMME contain genes, reactions, and metabolites that are not included in models produced by other approaches, since these two algorithms produce much larger models. Interestingly, those components tend to be shared by the two techniques, highlighting a similarity in the algorithms’ methods of pruning of the base model.

**Figure 2:**
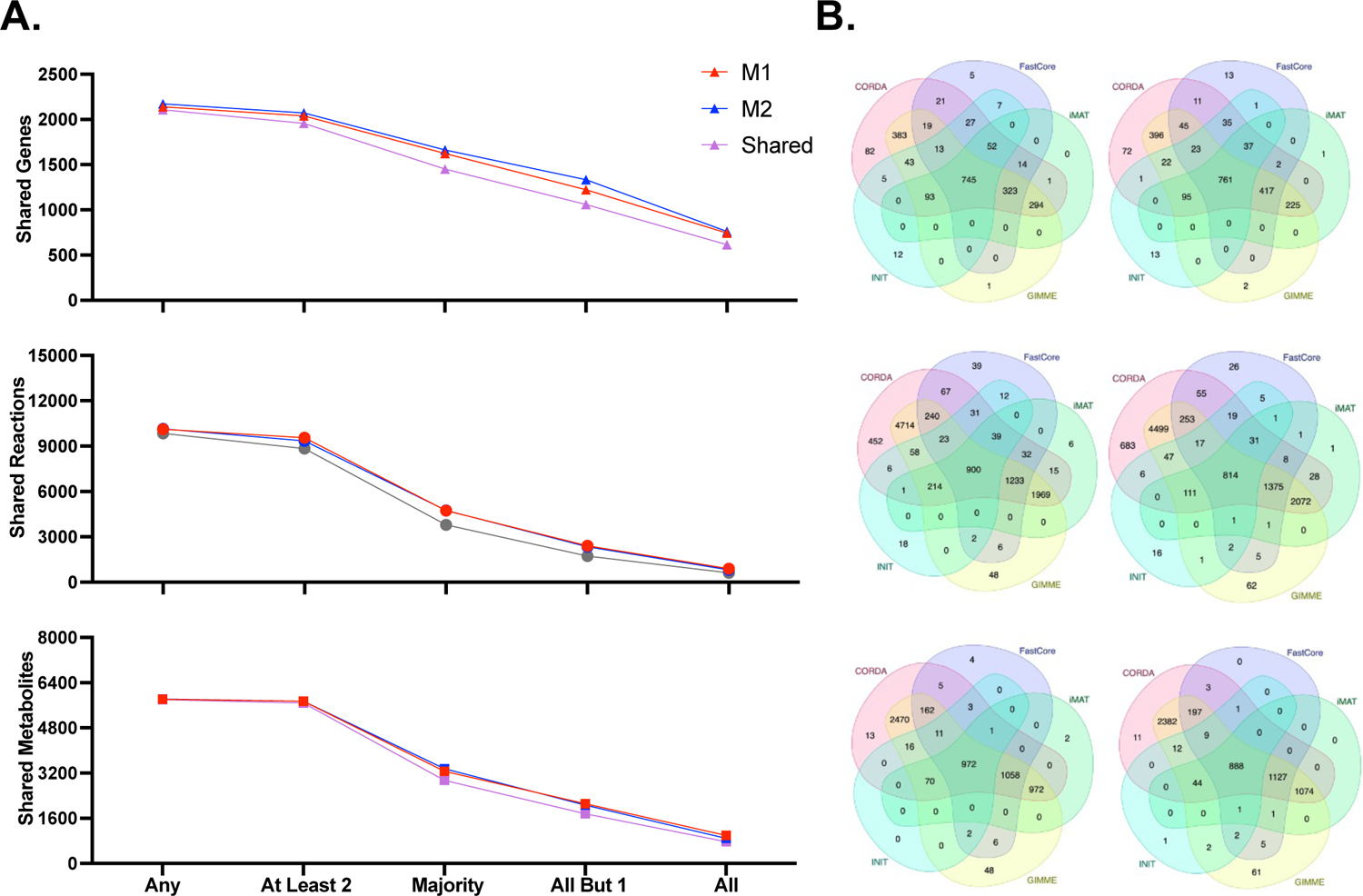
Comparison of model ensembles. **A.** Number of genes (top), reactions (middle), and metabolites (bottom) that are found in the GEMs for various ensemble thresholds. **B.** Venn diagram showing the number of model components shared between models generated by each integration algorithm: genes (top), reactions (middle) and metabolites (bottom) for M1 (left) and M2 (right).

### 4.2 Consolidated M1- and M2-specific models

Having demonstrated the quality of individual models, we aimed to arrive at a single consensus model for each of the M1 and M2 macrophage subtypes, to clearly understand metabolic differences between the cells. For each subtype, we compiled a list of model components found in a majority (at least three) of the five generated models. That list of metabolic components was then used to develop consensus M1- and M2-specific models. We applied the RedHuman technique to develop the minimal models connecting the model components consistently found in the M1 and M2 model ensembles. The RedHuman pipeline uses a human base model (Recon3D), thermodynamic information on human metabolic reactions, and a set of desired model components to generate high confidence reduced or core models. We therefore generated consensus models characterizing the M1- and M2-specific metabolism. The two consensus models are of similar size (**Figure 3A**) and share most model components, highlighting the overall similarity within the macrophage cell type. However, differences that arise between the two phenotypes are related to subtype-specific function.

**Figure 3:**
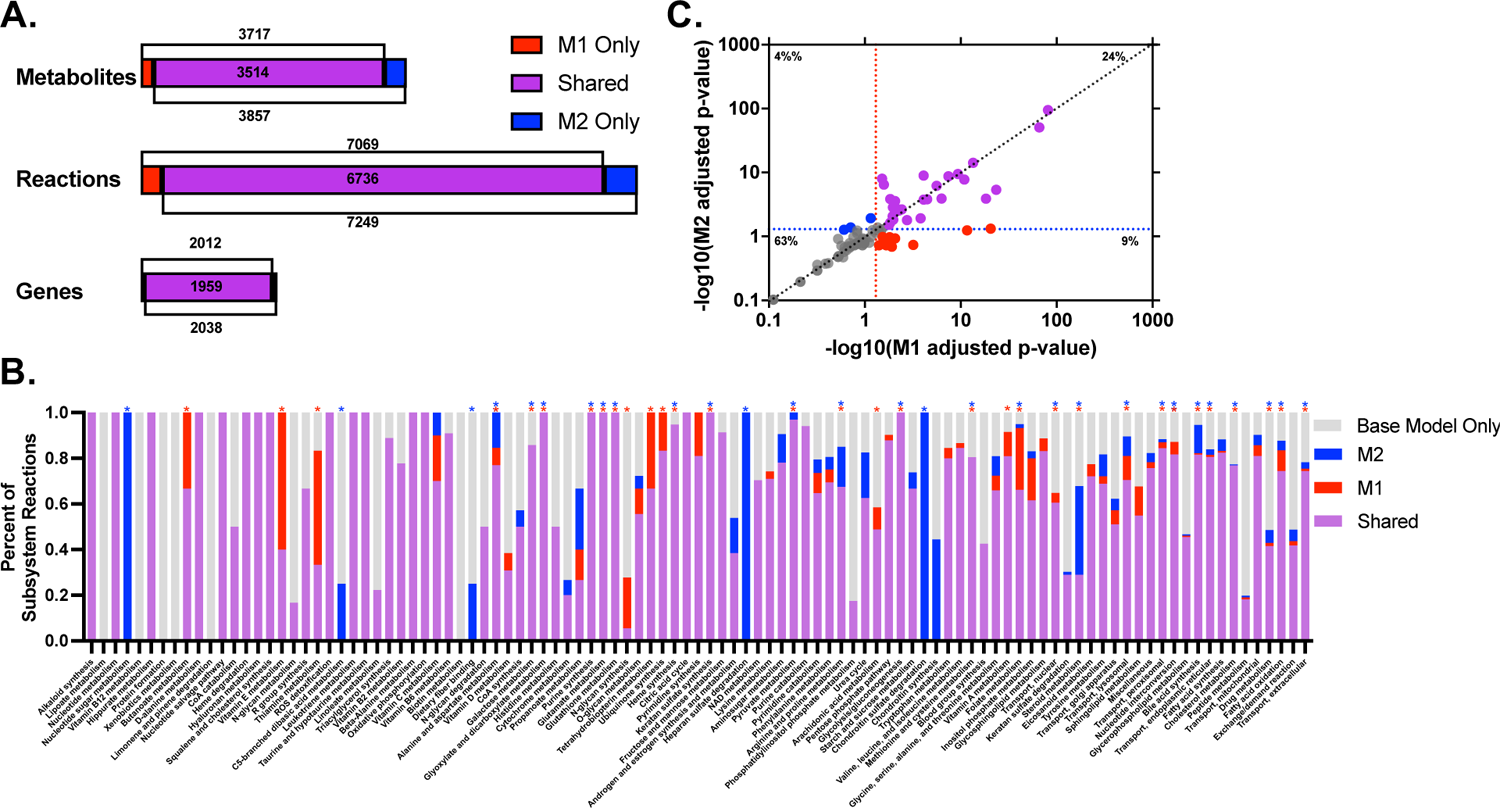
Characteristics of the consensus models. **A.** Sizes of M1- and M2-specific models, including shared (purple) genes, reactions, and metabolites, and components only found in the M1 (red) or M2 (blue) models. **B.** Comparison of the pathway composition for M1 and M2 models, relative to the pathway size in Recon3D. The pathways significantly enriched in M1 and M2 are marked with a red or blue asterisk, respectively. **C.** Results from FEA for the presence or absence of M1 and M2 metabolic pathways. Red and blue dotted lines represent significant values of *p*=0.05 for M1 and M2 models, respectively; the diagonal line represents the case where M1 and M2 subsystems are equally enriched.

Network composition is known to be a major cause of variation in predicted cell activity^67^. To assess structural differences between the macrophage subtypes, we analyzed the presence of reactions in specific metabolic pathways in the M1 and M2 models, relative to the Recon3D base model (**Figure 3B**). Some metabolic pathways (including lipoate metabolism, vitamin B12 metabolism, protein formation, xenobiotics metabolism, and biotin metabolism) were not found to be present in either macrophage subtype. Certain complete pathways (nucleotide sugar metabolism, heparan sulfate degradation and chondroitin sulfate degradation) are predicted to be found only in the M2 subtype. Portions of other pathways are only found in the M2 subtype (nucleotide sugar metabolism, C5-branched dibasic acid metabolism, and dietary fiber binding). Finally, there are subsystems that are more prevalent in M2, compared to the M1 state, including propanoate metabolism, the urea cycle, and alanine and aspartate metabolism.

Interestingly, there are no complete pathways found exclusively in the M1 phenotype. However, several pathways have reactions that are found only in the M1 model. This includes butanoate metabolism, vitamin E metabolism, thiamine metabolism, vitamin D metabolism, N-glycan synthesis, ubiquinone synthesis, pyrimidine metabolism, and arachidonic acid metabolism. Additionally, certain pathways contain more reactions present in M1 than in M2 (such as the xenobiotics pathway, the metabolism of Vitamin D, fatty acid oxidation, and folate metabolism).

To more robustly quantify differences between the M1 and M2 consensus models, we performed flux enrichment analysis (FEA)^41, 68^. The approach provides a statistical metric for the overrepresentation of metabolic components in a model relative to a reference state. We compared each subtypes’ reactions to the Recon3D base model and calculated the enrichment score and *p*-value for each metabolic pathway. The results of the approach are shown in **Figure 3C**, where the adjusted p-value for M1 and M2 pathways are shown on the *x*- and *y*-axes, respectively. Dots to the right of the vertical *p*=0.05 dashed red line on the *x*-axis are pathways considered highly enriched in the M1 model, and dots above the horizontal *p*=0.05 dashed blue line are highly enriched in the M2 model. The list of pathways is provided in **Supplementary Table 2**.

Most metabolic pathways that are enriched compared to the Recon3D base model are present in both macrophage subtype models (highlighted in purple), comprising 24% of all metabolic pathways. This implies those metabolic pathways are particularly important to macrophages but are not subtype-specific. The pathways include glycerophospholipid metabolism, peptide metabolism, N-glycan metabolism, and pyruvate metabolism, all of which are traditionally outside of standard metabolomic analyses but may be of particular importance to macrophage function.

M1-specific enriched pathways (9% of all metabolic pathways) include pyrimidine metabolism and propanoate metabolism, which have both been heavily implicated with the M1 macrophage immune response^69–72^. Additional pathways include the PPP, which is viewed as indicative of metabolic reprogramming towards an M1 state, along with ubiquinone metabolism and the urea cycle^26, 73^.

M2-specific significant pathways, which constitute 4% of all pathways consist of aminosugar metabolism, C5-branched dibasic acid metabolism, eicosanoid metabolism, and nuclear transport. In particular, the metabolism of eicosapentaenoic acid-derived eicosanoids has previously been determined to be a major sign of macrophage polarization towards an M2 state^74^.

### 4.3 Analysis of Model Flux Distributions

A cell’s distribution of metabolic fluxes is a useful metric for evaluating cellular state and comparing between distinct phenotypes. We therefore compared the predicted metabolic flux distributions for the two macrophage subtypes in order to determine reactions and pathways that are differentially utilized. Namely, we performed flux sampling with the RHMC algorithm, generating 50,000 flux distributions for each model. We then performed multi-objective FBA to predict the optimal flux distribution with the chosen objectives. By performing a comparative analysis on those predicted fluxes for shared reactions, it is possible to characterize the metabolism of the two subtypes.

With sampling, we found that most metabolic reactions show nearly equivalent mean flux values in the M1 and M2 subgroups, as shown in the diagonal trend seen in **Figure S1A**. Similarly, reactions exhibiting small or wide variance in metabolic flux tend to do so for both M1 and M2, as shown in the roughly diagonal trend seen in the standard deviation plotted in **Figure S1B**. However, the off-diagonal points (where flux values differ substantially for the M1 and M2 case) indicate differential fluxes between the two cell subtypes. When using MOFA optimization (**Figure S1C**), we see a near-inversion of the metabolic state, with many reactions’ fluxes showing different directionality and a minimal number along the diagonal. This suggests that, if we assume maximal orientation of cellular material towards divergent cellular goals (through selection of objective functions), we will see vastly distinct metabolic states.

To better capture the differences between sampled fluxes, we used the Kullback-Leibler (KL) divergence metric to compare the flux distributions of metabolic reactions present in both the M1 and M2 consensus models. The KL divergence metric categorizes the dissimilarity between two distributions. Thus, we performed a pairwise analysis for each reaction shared between the two models. As shown in **Figure 4A**, the flux distributions of only 19% of the reactions present in both the M1 and M2 consensus models are in close agreement, with “low” divergence (KL divergence value < 0.05). Approximately 20% of the shared reactions have very different flux distributions or “high” divergence in the M1 and M2 models (KL divergence value > 0.5) groups. The majority of shared reactions (61%) had “medium” divergence across the two macrophage subtypes.

**Figure 4:**
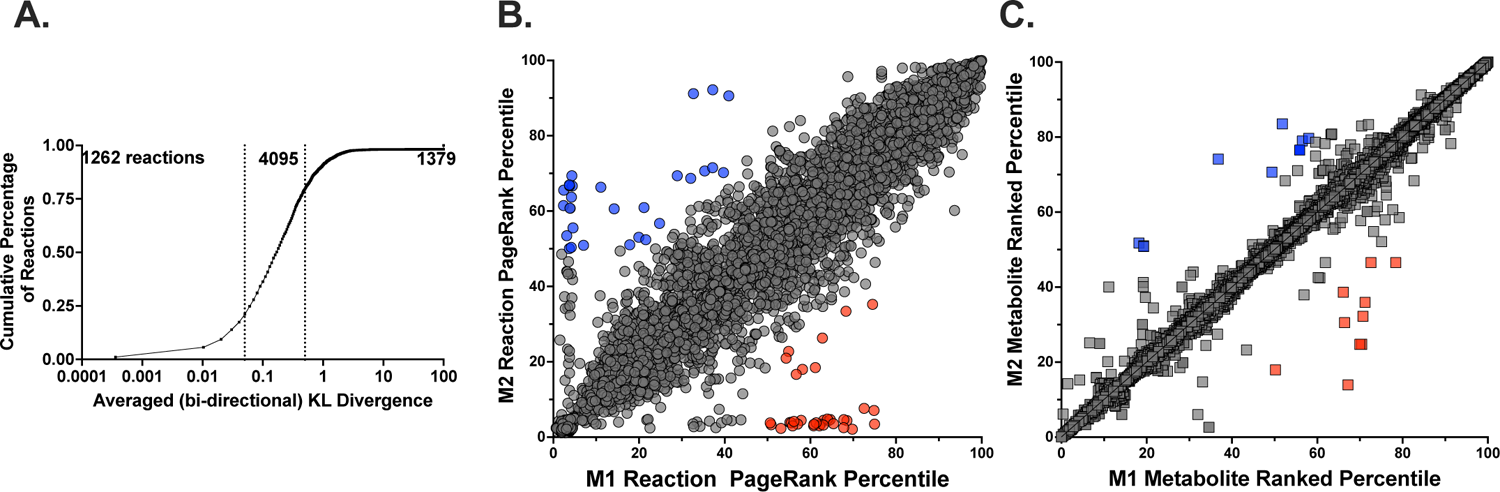
Model flux predictions. **A.** KL divergence values from pairwise comparison of flux distributions for reactions present in the M1 and M2 consensus models. **B.** Comparison of the weighted PageRank scores for M1 and M2 consensus models for all shared metabolic reactions (circles). *Red*, reactions that were highly important in M1 but not M2; *blue*, reactions that were highly important in M2 but not M1. **C.** Comparison of the metabolite rankings between the M1 and M2 consensus models. Subtype-specific metabolites: *red*, M1 and *blue*, M2.

At the pathway level, there are key differences in the flux distributions between the M1 and M2 consensus models. We calculated the average flux divergence score across the metabolic subsystems and found five pathways for which the reactions had significantly distinct flux distributions between the M1 and M2 models: butanoate metabolism, linoleate metabolism, alkaloid synthesis, nucleotide metabolism, and vitamin E metabolism. Each of those pathways has previously been shown to be involved in the anti-inflammatory or pro-inflammatory activity of macrophages^69, 75–80^. Thus, this quantitative analysis complements experimental evidence pointing to differences between macrophage subtypes.

We then calculated the relative importance of each metabolic reaction by pairing a graph theory-based calculation of network centrality (the PageRank algorithm) with the predicted metabolic fluxes. The flux-weighted centrality scores were compared between the two subtypes and are shown in **Figure 4B**. Thirty-three reactions emerged as only important for M1 cells, and 28 emerged as only important for M2. Most of those were transport reactions, highlighting variation in preferred metabolic fuel sources. Specifically, import of alanine, L-glutamine, and His-Glue are more important for M1, while transport of L-leucine, chitobiose, and formate are more important for M2. Non-transport reactions that were impactful were generally related to central carbon metabolism and ganglioside metabolism for M1, with malic enzyme and ganglioside galactotransferase both emerging as important. M2 cells showed strong scores for Coenzyme-A-related reactions involved in fatty acid metabolism, with the production of both phytanyl-CoA and hexadecanoyl CoA found to be significant.

As a complement to the analysis of reaction importance, we sought to use a metabolite-centric approach to elucidate subtype-specific differences. We calculated the flux-sum values for each metabolite in the model, to understand the relative importance of each metabolite in the network. The flux-sum value is defined as one-half of the sum of fluxes in and out of a metabolite pool and is often used to characterize the metabolite’s turnover rate^81^. We compared the score for each metabolite present in the two cell type-specific metabolic models. This analysis revealed 19 metabolites whose usage differed substantially in the two subtypes: 9 specific to M1 and 10 specific to M2. Influential M1 metabolites include adenosine and G3P, which have both been connected to the immune response seen in macrophages, as well as several amino acids (arginyl-valyl-tryptophan and aspartyl-glutamate) and two coenzyme A-activated acyl groups ((7Z)-hexadecenoly CoA and (6Z,9Z)-octadecadienoyl CoA). M2 metabolites of importance include cholesterol ester, which has been suggested to be related to M2 polarization, D-mannose, which is thought to suppress inflammatory (M1) state, and four other coenzyme A-activated acyl groups (3-oxotridecanoyl CoA, (S)-3-hydroxytetradecanoyl CoA, (S)-3-hydroxyoctadecanoyl CoA, and nonanoyl CoA)^82–86^. The emergence of distinct CoA-related metabolites agrees with and builds upon past work emphasizing the divergent role of fatty acid metabolism in the M1/M2 paradigm^87–90^.

Altogether, by pairing predictions of metabolic flux distributions with network topology, and by evaluating metabolism from both a reaction- and metabolite-centric view, we produce an in-depth understanding of the divergent macrophage metabolic phenotypes.

### 4.4 Model validation

To validate the model predictions, we compared model composition and predicted flux with canonical characteristics of M1 and M2 macrophages, provided in **Table 1**. We identified nine metabolic features from the literature, and 78% of the commonly accepted differences between the M1 and M2 phenotypes are captured by our model predictions. This provides confidence in the model structure and flux distributions.

**Table 1.**
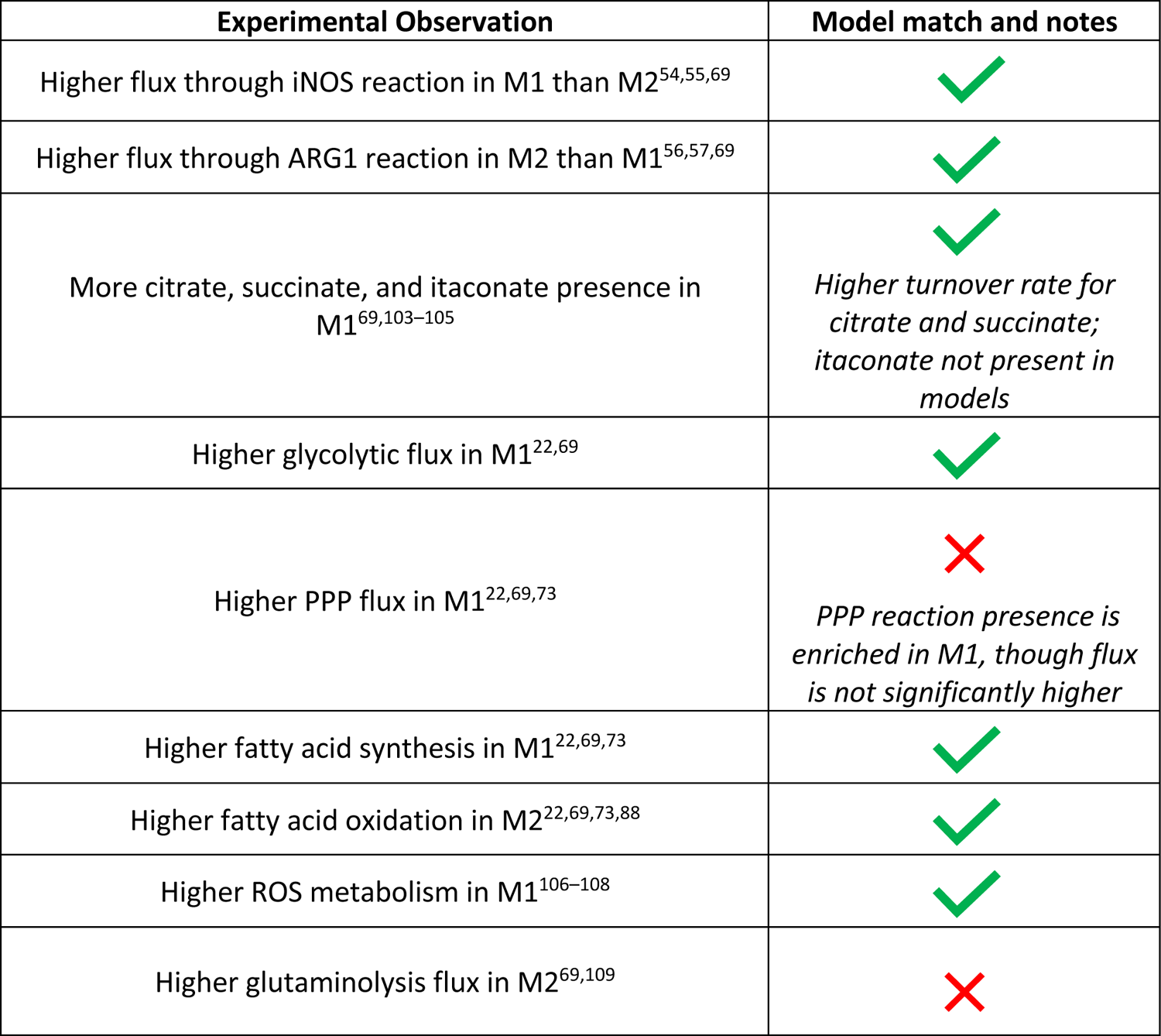
Qualitative comparison of model predictions and experimental observations

### 4.5 Analysis of model sensitivity to perturbation

Having constructed and analyzed genome-scale metabolic models for the M1 and M2 macrophage subtypes, we sought to identify metabolic perturbations that could push macrophages from a TAM-like state towards a pro-immune (and therefore, anti-cancer) condition. In order to accomplish this, we developed a sampling-based, objective function-independent algorithm modeled after the minimization of metabolic adjustment (MOMA) approach. This method identifies the minimal intervention needed to push the sampled flux distribution for a particular reaction in a candidate constraint-based model towards a desired flux state. More detail is provided in the *Methods and Materials* section.

Certain metabolic perturbations are predicted to alter the flux distributions of reactions that characterize the M2 metabolic phenotype. We first identified the top 10 M2-specific reactions (via the flux-weighted PageRank analysis) and performed individual and pairwise enzyme knockouts by systematically inhibiting flux through those reactions. We then sampled the knockout models and calculated the cosine distance between the sampled flux distribution of the knockout model and the baseline M2 model flux. This identified the flux distributions for reactions that are “highly divergent” based on the KL value between the knockout M2 model and baseline M2 model.

Interestingly, nearly all implemented perturbations caused the knockout M2 model to take on a phenotype that is distinct from the baseline M2 metabolic state (**Figure 5A**). This result suggests validity of using the flux-weighted centrality approach for finding potential metabolic targets. Furthermore, we observe clear benefit from the multi-target approach, as pairwise combinations tend to cause greater changes than single knockouts alone (compare off-diagonal and diagonal values in **Figure 5A**). The largest percent change for individual enzyme knockouts was achieved with the *glycine synthase* reaction, causing an 18% difference from the baseline M2 metabolic state, followed closely by *glyceraldehyde-3-phosphage dehydrogenase* and the formation of deoxy-fluvastatin, which each caused a 17% difference from the baseline M2 model^22, 91^. The greatest impact of pairwise knockouts was achieved when the phosphate-Na+ transporter (*Plt7*) and phosphatidylethanolamine N-methyltransferase (*PETOHMr_hs*) reactions were shut off in concert, which caused a 30% divergence away from the baseline M2 state.

**Figure 5:**
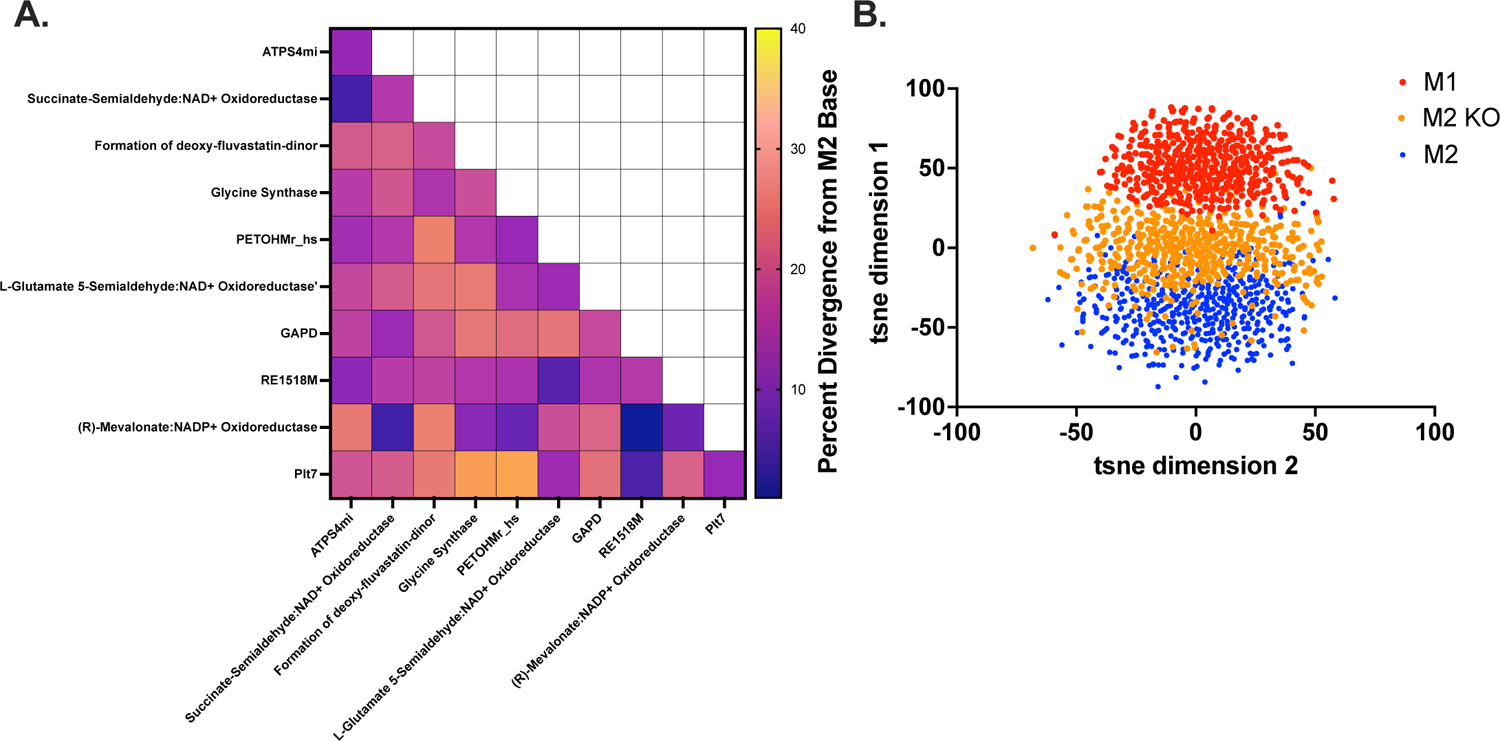
Reaction knockout analysis. **A.** KL divergence for M2 fluxes in the baseline model and knockout model for all highly divergent reactions identified by the KL metric shown in Figure 4A. **B.** Visualization of metabolic flux samples in low-dimensional space for the baseline M2 model (*blue*), knockout M2 model (*orange*), and baseline M1 model (*red*).

Notably, that pairwise effect was slightly synergistic, as the combination outcome was greater than the additive effect of each knockout individually (11% and 17%, respectively). The second-largest intervention was pairing *glycine synthase* with the *Plt7* reaction, which produced a 29% difference between the knockout and baseline M2 models.

Inhibiting metabolic reactions in the M2-specific model is predicted to move the flux distribution away from the baseline M2 metabolic state. After performing the enzyme knockouts, we performed dimensional reduction using t-SNE on the 50,000 sampled flux distributions for the M2 model for which *Plt7* and *PETOHMr_hs* were inhibited. **Figure 5B** shows 1,000 of the sampled flux distributions, for each of the three conditions (baseline M1, baseline M2, and *Plt7/PETOHMr_hs* combination knockout M2). Each point in the figure is a flux distribution vector for all reactions shared between the metabolic models, represented in low-dimensional space. This analysis shows that the metabolic states of the knockout model are positioned between the M1 and M2 baseline models, showing the metabolic perturbation elicits a shift in the metabolic phenotype towards the M1 metabolic state.

Overall, we identify metabolic reactions that define the M2 metabolic state and predict how targeting those reactions influences the shift towards a more M1-like metabolic phenotype.

## 5 Discussion

Researchers have long acknowledged the importance of the immune system in cancer and have appreciated the influence of intracellular metabolism on observed cellular activity. However, cancer immunometabolism, the convergence of immunology and metabolism in cancer, has only seen intense interest in recent years. It is thought that a proper understanding of the metabolic mechanisms impacting immune cell behavior may inform promising therapies in cancer^92, 93^. Excitingly, the increased interest in cancer immunometabolism has coincided with a significant expansion in our technical ability to measure cell content, with large-omics datasets generated by high-throughput experimental approaches^94^. Those datasets provide substantial insight into the variation between cells and the effect of the environment on observed phenotype^95–97^. However, a complete characterization of metabolism can be difficult for many reasons, including the limited availability of a sufficient number of cells and the dynamic nature of metabolism. Genome-scale modeling of metabolism has emerged as a possible solution to comprehensively understand cellular metabolism, allowing integration of -omics datasets to generate novel biological insight. The models integrate and synthesize data to mechanistically understand metabolism.

In this work, we generate M1- and M2-specific GEMs of human macrophage metabolism. The models are the first to study human macrophage subtypes in cancer at the genome-scale and are built specifically with patient-derived transcriptomics data. The models are consistent with existing knowledge of cancer-associated macrophages and predict clear differences between the metabolism of the M1 and M2 subtypes, both structurally (the composition of the metabolic network) and functionally (utilization of the metabolic reactions).

In making M1- and M2-specific metabolic models, we combine existing approaches and algorithms into a coherent methodology that can be applied to other cell types and conditions in the future. In particular, by generating a set of models using a variety of model pruning algorithms, we developed an approach that limits bias resulting from the selection of a single data-integration technique. The models were analyzed as an ensemble, an approach that has previously been shown to increase model predictive accuracy, but has not been applied to sets of models generated from distinct -omics data integration techniques^98–100^. The large variation within the ensemble suggests that future work should identify additional methods to remove user-imposed bias in model generation, particularly as the availability of -omics data increases. For both the M1 and M2 subtypes, we consolidated the ensemble of models into a single consensus model for each subtype. We assessed model composition and combined a variety of analytical and statistical methods to characterize the divergence and similarities of the two macrophage subtypes. The approach also allows for rapid assessment of interventions to alter the predicted metabolic state. The predictions reveal promising strategies that can be experimentally tested in future work.

With established and validated GEMs of M1 and M2 macrophage metabolism, there are many newly available directions of research. For example, in this work we operate with the standard FBA assumption that the system is at a steady state. Future efforts could relax that constraint, thereby using these models to predict dynamics and the way in which cellular metabolism changes over time^101^. Second, though the M1-M2 scheme is relatively well validated and supported in previous studies, it is a substantial oversimplification of the nature of macrophage phenotypes. Namely, macrophages are extremely plastic, and demonstrate significant heterogeneity, both across a population and throughout a single cell’s lifespan. Future work may assess and quantify macrophage metabolic heterogeneity. Finally, these cells do not exist in isolation, but are influenced by and affect neighboring cells. There has been work to understand metabolic interactions between GEMs, but it has largely been limited to microbial interactions. Future studies may apply those techniques to understanding the tumor microenvironment as well, including GEMs of immune cells such as the ones produced in this work^102^.

In summary, we utilized genome-scale metabolic modeling to investigate the differences in M1 and M2 macrophage subtypes in colorectal cancer. We compared model composition and predicted metabolic activity, furthering our understanding of the subtypes’ metabolic states and capabilities. Furthermore, we identified potential metabolic targets that might push M2 macrophages toward an anti-cancer condition, potentially improving patient outcomes. Overall, this work is a substantial advance in our understanding of macrophage metabolic state and activity.

## 6 Limitations of the study

We acknowledge some limitations of our work. We analyzed and simulated the consensus M1 and M2 models, comparing model components and assessing their network flux distributions. We found clear and consistent metabolic signatures particular to each phenotype and find literature support for many of the model predictions. However, because the models are built directly from patient data, the source or exact cause of those observed signatures cannot fully be determined. We attempted to limit the effect of patient-to-patient heterogeneity by merging the measurements into a pseudo-bulk dataset, but it is not entirely clear how much of the metabolic phenotype we predict is due to differences in the metabolic states of M1 and M2 macrophages, patient-specific differences, or metabolic reprogramming caused by cancer cells and other cells in the tumor microenvironment. Similarly, due to a paucity of experimental data characterizing the intracellular metabolism of macrophages (outside of major central metabolic pathways), it is difficult to make direct comparisons that fully validate our predictions.

Nevertheless, the experimental observations collected in Table 1 provide the most relevant and informative insight for model validation. Future work can be done to test the model predictions, starting with *in vitro* experiments and progressing to *in vivo* studies.

## Acknowledgements

This work was supported by the National Cancer Institute of the National Institutes of Health grant 1U01CA232137 and administrative supplement (to S.D.F.). The authors thank members of the Finley research group for critical feedback and Drs. Vassily Hatzimanikatis and Maria Masid for their input and assistance with the RedHuman tool.

## 7 Author contributions

P.E.G. and S.D.F. conceived of the presented idea. P.E.G. planned and carried out computational model development and simulations. S.D.F. supervised the project and provided financial support. All authors discussed the results and contributed to the final manuscript.

## Star Methods

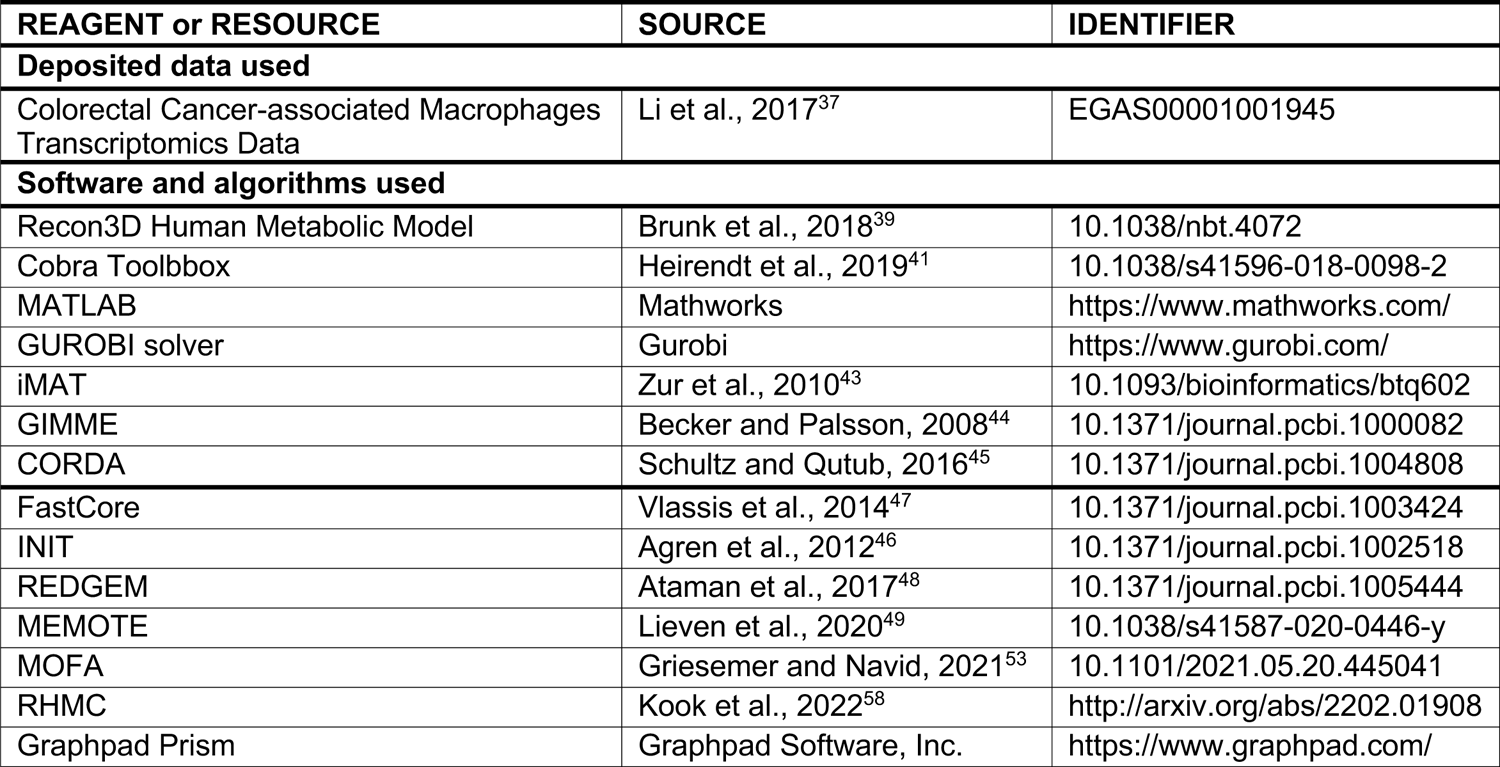

